# Exploring the Limits of EPR-driven Tumor Accumulation with Non-opsonizing Nanomaterials

**DOI:** 10.1101/2024.05.07.592926

**Authors:** Rinalds Serzants, Baiba Svalbe, Irina Cesnokova, Gundega Stelfa, Antons Sizovs

## Abstract

The Enhanced Permeability and Retention (EPR) effect is a foundational concept used to rationalize nanomedicine development for cancer treatment and diagnostics. The attainable efficacy of passive tumor targeting due to EPR remains ambiguous owing to pervasive opsonization of nanoparticles. To address this, we developed nanomaterials with complete resistance to opsonization, exceptionally long systemic circulation, and used them to study the limits of the EPR in triple-negative breast cancer. Tumors exerted no impact on pharmacokinetic profiles, which were indistinguishable between healthy and tumor-bearing mice. Tumors were the primary accumulation sites and our data revealed that the maximum average achievable tumor accumulation via EPR is proximate to 60 %ID/g, tumor-to-liver selectivity is 4-to-1, and the optimal D_H_ to fully exploit EPR lies between 18 and 54 nm. The significant heterogeneity observed in tumor accumulation, however, indicates that nanomedicines cannot achieve consistent efficacy across different patients by relying solely on EPR.

---

In 1984, Maeda and colleagues reviewed and discussed the mounting evidence for high tumoritropicity displayed by macromolecules and protein-polymer conjugates^1^. Two years later, in 1986, they published a seminal paper describing a new concept for macromolecular accumulation in tumors, which now is known as Enhanced Permeability and Retention (EPR) effect^2^. The EPR has become the major phenomenon to rationalize development of macromolecular/nano-sized anticancer therapeutic and diagnostic agents^3^. Recently, mechanistic aspects of the nanomedicine accumulation in solid tumors have been closely investigated, uncovering that in addition to leaky vasculature and reduced lymphatic drainage in tumors, accumulation results from transcytosis^4,5^. While they were excellent in shedding light on contribution of endothelial translocation, little attention was paid to opsonization of nanomaterials used in these studies. It is well understood that to maximize the exploitation of EPR, nanomedicines require prolonged systemic circulation to effectively reach tumor site^6^. This has been addressed by clever strategies, like overwhelming Kupffer cells uptake capacity thus mitigating accumulation in liver^7^, but is more commonly approached by passivation of nanoparticles to reduce non-specific opsonization^8,9^. Opsonization and protein corona formation renders nanomedicines recognizable by mononuclear phagocytic system (MPS)^10–12^. This not only leads to rapid clearance, but also unpredictably alters the properties of nanomaterials, importantly – the size. Consequently, while numerous studies have demonstrated nanoparticle accumulation in tumor and arrived at conclusions regarding the influence of nanomaterial size on passive tumor accumulation, a definitive consensus on both the optimal size for nanomedicines and the maximum attainable accumulation in tumors by EPR has yet to be established (see Kang et al. for discussion^13^). In this study, we engineered non-opsonizing nanomaterials optimized for prolonged circulation. Utilizing their size-driven biodistribution, we determined the EPR-driven accumulation efficiency in 4T1 model of TNBC tumors in immunocompetent mice, offering a refined understanding of EPR potential and limits in TNBC tumor and offering a tool to study EPR in other cancer models.

## Non-opsonizing nanoparticles

To engineer non-opsonizing nanoparticles we looked to PEG, a polymer known to provide resistance to opsonization and extend nanoparticle circulation^9,14^. Complete PEGylation of the nanoparticle surface, however, is practically unattainable leaving the potential sites for protein adherence^15^. Therefore, to take the full advantage of non-opsonized properties of PEG, we devised a unimolecular surrogate for nanoparticles – bottle brush polymers that almost entirely are composed of PEG. To create brush polymers with significant volumetric prominence we used a large, 2000 Da, PEG monomer that has 45 repeat units (α-methoxy-Ω-methacrylamide polyethylene glycol, 2kPEG-NMA). We chose methacrylamide derivatives to generate brush polymers resistant to hydrolysis. A 2 mol% of aminoethylmethacrylamide was copolymerized with the PEG monomer to serve as a chemical handle for incorporation of fluorescent dyes (**Fig. 1a**). We analyzed prepared Brushes with dynamic light scattering (DLS), chose three brush polymers that cover most interesting 15-50 nm size range ^13,16,17^, and labeled them with AlexaFluor dyes. Namely, for this study we selected three brush polymers and assigned names according to their hydrodynamic sizes: Brush-18 (D_H_=18 nm), Brush-29 (D_H_=29 nm), Brush-54 (D_H_=54 nm) (**Fig. 1b**).

**Fig. 1.**
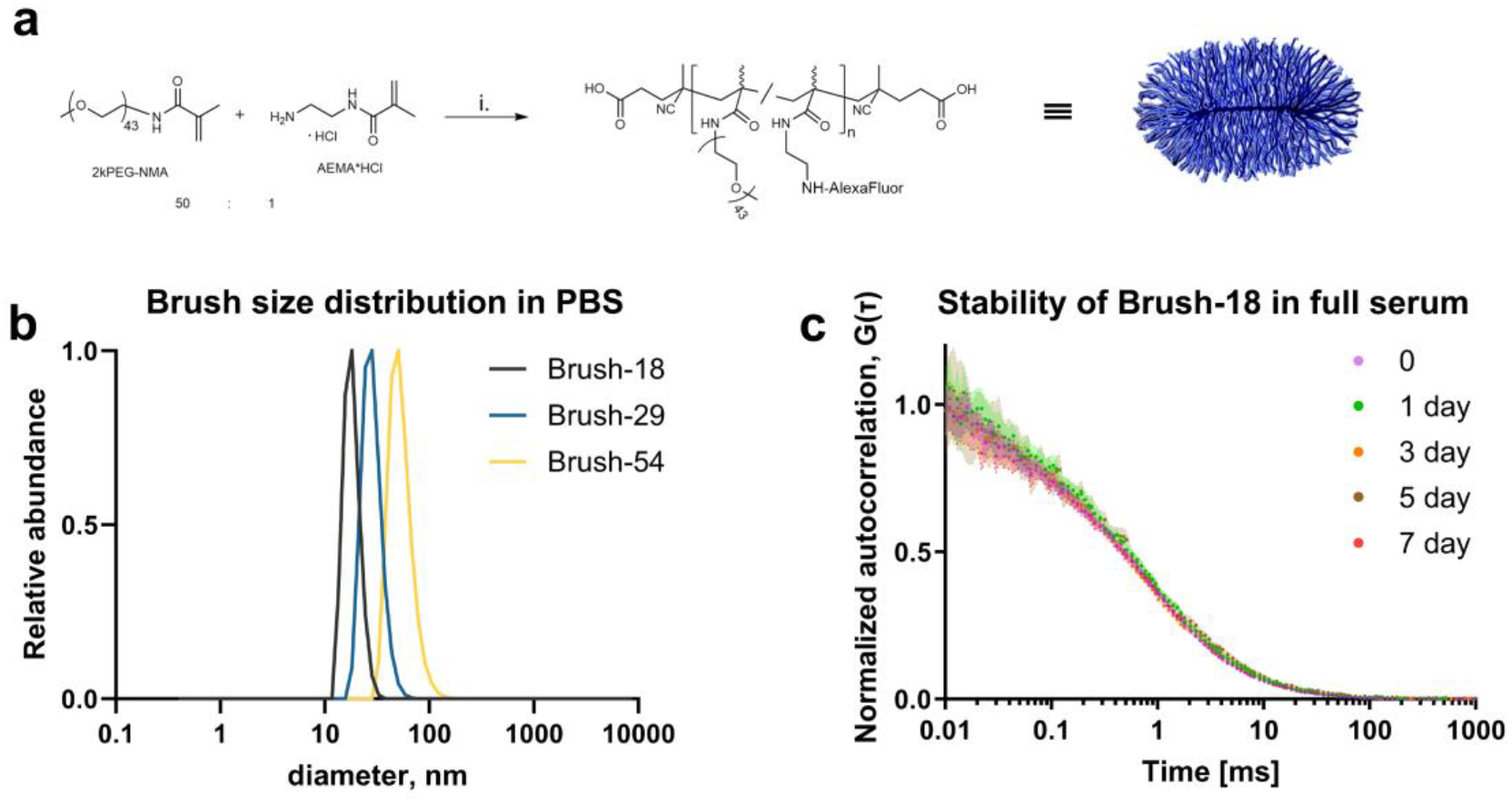
Bottle brush synthesis and characterization. **a**, Syntheses were carried out by (i) RAFT polymerization of 2kPEG-NMA and AEMA monomers (50 to 1 mol ratio), removal of the trithiocarbonate group, followed by conjugation of AlexaFluor dyes and caping of any leftover primary amines with mPEG_12_-NHS. **b**. Dynamic light scattering (DLS) analysis of three polymers used for further investigation had D_H_=18 nm, 29 nm, 54 nm. **c**. Fluorescence correlation spectroscopy (FCS) autocorrelation curves did not change during the 7 day period of incubation in full mouse serum at 37 °C, indicating there were no changes in Brush diffusion and no opsonization or protein corona formation.

Upon entering the circulatory system, nanoparticles are rapidly opsonized, leading to alterations in their physicochemical properties, most notably their size. We employed fluorescence correlation spectroscopy (FCS) to demonstrate resistance of prepared Brushes to non-specific opsonization by serum proteins. FCS is conceptually similar to DLS as both techniques rely on statistical correlations (autocorrelation) in the time domain, but FCS uses fluctuation of fluorescence intensity and therefore can be used to investigate fluorescent materials even in the presence of proteins in high concentrations^18^. To closely mimic *in vivo* conditions, we incubated Brushes in full mouse serum at 37 °C. Albumin is the most abundant serum protein and has D_H_ of ∼8.5 nm. Adsorption of albumin (or other serum proteins) to our nanomaterials would lead to a significant increase in size and result in a profound shift of Brush autocorrelation curves, *G*(τ)^19^. Our data, however, revealed that even after one week of incubation, autocorrelation curves of Brushes did not change (**Fig. 1c and Supplementary Fig. 1**). This indicates that these materials are efficient in resisting the non-specific opsonization and do not acquire a protein corona. Furthermore, the stability of the autocorrelation curves implies that Brush polymers neither degrade nor aggregate, ensuring their consistent size in circulation.

## Pharmacokinetics and biodistribution in healthy mice

Before exploring the EPR of tumors, we assessed the PK and BD of engineered non-opsonizing nanomaterials in healthy immunocompetent mice, to establish a baseline, and to confirm that their non-opsonizing properties can result in extended circulation times, avoid rapid clearance by MPS and lead to broad biodistribution. To achieve this, we administered equal amounts of Brush-18, Brush-29, and Brush-54 to Balb/c mice using both intravenous (i.v.) and intraperitoneal (i.p.) routes.

Following both administration routes, Brush polymers exhibited remarkably long circulation times (**Fig. 2a**). Thus, the longest-circulating Brush-18 was present in blood at 4.9±0.3 %ID/mL concentration at 14 days after *i*.*v*. administration. A two-compartment pharmacokinetic model was utilized to fit the measured concentrations, revealing elimination half-lives (t_1/2_β) of 3.5, 2.7, and 1.2 days for Brush-18, Brush-29, and Brush-54 respectively, with corresponding mean residence times (MRTs) of 5.00, 3.67, and 1.72 days after *i*.*v*. administrations (**Fig. 2b**). Administration to peritoneal cavity sidesteps the initial high-concentration ‘plug’ that results from *i*.*v*. bolus administration. It avoids possible oversaturation of MPS (Kupffer cells)^7^, allowing for PK/BD profile to be primarily determined by the nanomaterial size. This route, however, requires the ability to escape peritoneal cavity, which primarily drains by lymphatic system^20^. All three Brushes exited peritoneum with indistinguishable efficiency and the highest blood concentrations were observed at 8 hours post administration (42.4±3.5, 35.9±2.5 and 43.2±1.7 %ID/mL for Brush-18, Brush-29, and Brush-54, mean±SEM). Clearance half-lives remained very long (**Fig. 2b**), confirming that these materials are well suited to reach tumor via systemic circulation and exploit the EPR. Rather counterintuitively, circulation was inversely dependent on Brush size (Fig. 2d), suggesting that ability to penetrate through the tissues rather than extravasation was the determining factor.

**Fig. 2.**
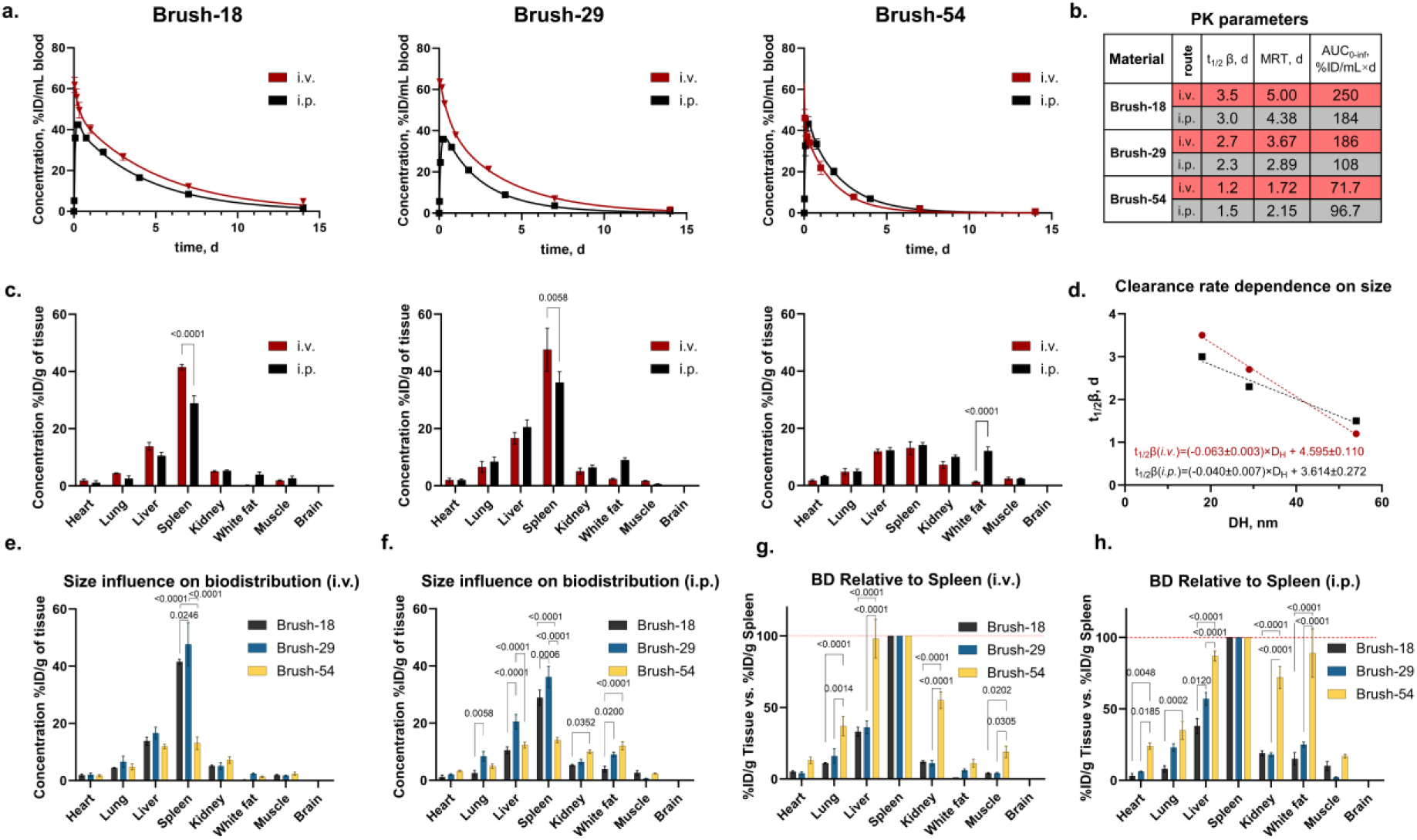
Pharmacokinetics and biodistribution in healthy immunocompetent mice. **a**, Pharmacokinetic profiles of Brush-18, Brush-29 and Brush-54 following i.v. (red line and symbols) and i.p. (black line and symbols) administration routes (n=4-6 mice, averages ± SD). **b**, Pharmacokinetic parameters obtained by fitting blood concentration data into two-compartment (i.v.) and extravascular two-compartment (i.p.) pharmacokinetic models: clearance half-life (t_1/2_β), mean residence time (MRT), Area Under the Curve from time zero to infinity (AUC_0-inf_). **c**, Biodistribution profiles of Brush-18, Brush-29 and Brush-54, 2 weeks after i.v. (red bars) and i.p. (black bars) administration routes. **d, e**, Influence of Brush size on biodistribution profiles two weeks after i.v. administration. **f**, Influence of Brush size on biodistribution profiles two weeks after i.p. administration. **g**, Influence of Brush size on accumulation in tissues versus accumulation in spleen two weeks after i.v. administration. Values are averages of relative tissue concentrations calculated individually for each mouse. **h**, Influence of Brush size on accumulation in tissues versus accumulation in spleen two weeks after i.p. administration. Values are averages of relative tissue concentrations calculated individually for each mouse. For **c, e**-**h**, n=4-5 mice, average ± SEM).

We assessed the biodistribution by measuring Brush concentrations in tissue homogenates two weeks after administration. All groups displayed broad biodistribution (**Fig. 2c**). As expected, organs characterized by the presence of large “exits”, such as 250-1200 nm-wide slits in spleen^21^ and 50-300 nm fenestrations in sinusoidal endothelial cells of liver^22^, had the highest Brush concentrations. Biodistribution profiles between *i*.*v*. and *i*.*p*. routes were remarkably similar, with measurable differences only in spleen and, surprisingly - white fat. Although these differences were very moderate, they reaffirmed the concern stated above that during bolus *i*.*v*. administration, the initial high concentration can influence the biodistribution profile.

Size of the Brush had little influence on BD following *i*.*v*. administration, with spleen being a notable exception (**Fig. 2e**), but it had a noticeable impact in case of i.p. (**Fig. 2f**). Brush-29 showed the highest accumulation in nano-penetrable organs, i.e. spleen (for both i.v. and i.p. routes) and liver (i.p. route). We also examined the biodistribution selectivity based on ease for extravasation, by normalizing concentrations in tissues against the corresponding concentrations in spleen of the same animal (**Figs. 2g and 2h**). Following *i*.*v*. and more significantly – *i*.*p*. administration routes, the relative accumulation of Brushes in organs with less permeable vasculature increased with size. These findings corroborate the conclusions based on differences in PK profiles, and emphasize that while passive extravasation is crucial (the highest Brush concentrations observed in the spleen, followed by the liver), it is the capacity of a nanomaterial to diffuse through interstitial spaces and subsequently re-enter circulation that ultimately determines distribution to less permeable tissues.

## Pharmacokinetics and biodistribution in TNBC tumor-bearing mice

To investigate the EPR effect and its limitations we used orthotopic 4T1 triple-negative breast cancer model in Balb/c mice. Consistent with our approach in evaluating PK and BD in healthy animals, we opted for immunocompetent mice to avoid ambiguity inherent to cancer models in immunocompromised hosts. 4T1 model is phenotypically similar to and shares substantial molecular features with human TNBC^23–25^. Moreover, 4T1 is widely used TNBC model and is of particular interest to cancer research community.

Based on the findings in healthy mice, we chose intraperitoneal administration route. Despite activation of the innate immune system and macrophages in peritoneum^26,27^, all three Brushes effectively drained from the peritoneal cavity, navigated past the lymph nodes and entered circulation.

This serves as yet another evidence for the ability of developed Brushes to remain ‘invisible’ *in vivo* and explore EPR effect purely as a function of nanomaterial size. The pharmacokinetic profiles for each of the nanomaterials in the presence of TNBC tumor were only marginally different from the corresponding profiles in healthy mice (**Fig. 3a**). Furthermore, direct comparison of the measured Brush concentrations at matching time points revealed no statistically significant differences between the healthy and TNBC cohorts (**Supplementary Fig. 2**). This rejects the notion that tumor can act as a sink for nanoparticles that can draw them from the circulation due to the EPR effect. Pharmacokinetic profiles of Brush-18, Brush-29, and Brush-54 reflected a counterintuitive trend previously observed in healthy mice and circulation exhibited an inverse relationship with Brush size (**Fig. 3b**).

**Fig. 3.**
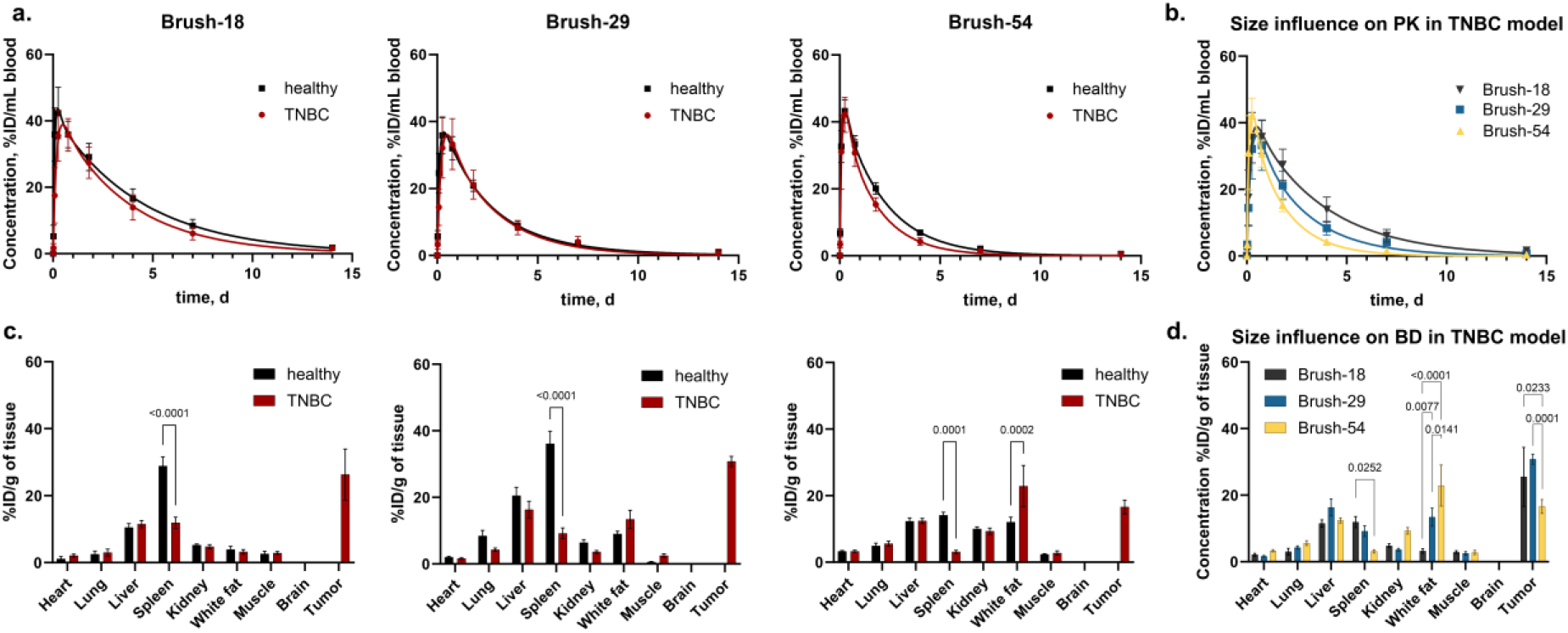
Pharmacokinetics and biodistribution in 4T1 TNBC tumor-bearing mice. **a**, Comparison of pharmacokinetic profiles of Brush-18, Brush-29 and Brush-54 in healthy (black symbols and lines) and TNBC mice (red symbols and lines) following intraperitoneal administration (n=4-6 mice, average± SD). **b**, Comparison of pharmacokinetic profiles between Brush-18, Brush-29 and Brush-54 in TNBC mice (n=4-6 mice, average± SD). **c**, Comparison of biodistribution profiles for Brush-18, Brush-29 and Brush-54 in healthy (black symbols and lines) and TNBC mice (red symbols and lines) 2 weeks after intraperitoneal administration (n=5 mice, average± SEM)**d**, Influence of Brush size on biodistribution profiles 2 weeks after intraperitoneal administration (n=5 mice, average± SEM).

The presence of a TNBC tumor did not markedly alter the biodistribution profiles, with the spleen being a significant exception (**Fig. 3c**). Profound splenomegaly is characteristic of 4T1 TNBC model and is associated with hematopoietic stem and progenitor cell infiltration and hematopoiesis that is skewed to myelopoiesis^28^. Large number of myeloid-derived suppressor cells in spleen are capable of engulfing nano-sized objects, however we observed significant reduction in splenic accumulation of Brushes. In comparison to the Brush concentrations in spleens of healthy mice, the concentrations in TNBC groups decreased 2.4, 3.9, and 4.5 times for Brush-18, Brush-29, and Brush-54, respectively. This effect correlated with the increase in Brush size and suggests that permeability of spleen vasculature in 4T1 tumor-bearing mice is greatly reduced.

Two weeks after administration, tumor was the clearly the major accumulation site in cases of Brush-18 and Brush-29, reaching 26.3±7.6 and 30.8±1.5 %ID/g of tumor, respectively (mean±SEM). Concentration of Brush-54, however, was 16.6±2.1 %ID/g and was not statistically different from Brush-54 concentrations in liver, kidneys and white fat (**Supplementary Fig. 3**). Moreover, both Brush-18 and Brush-29 more effectively accumulated in tumor than Brush-54 (**Fig. 3d**). We recognized, however, that biodistribution was complete for Brush-54 and nearly complete for Brush-18 and Brush-29 already at seven days after administration (**Fig. 3a**). During the subsequent period between days 7 and 14, tumors continued to grow, more than doubling in size (**Supplementary Fig. 4**). This growth effectively “diluted” the accumulated Brushes.

**Fig. 4.**
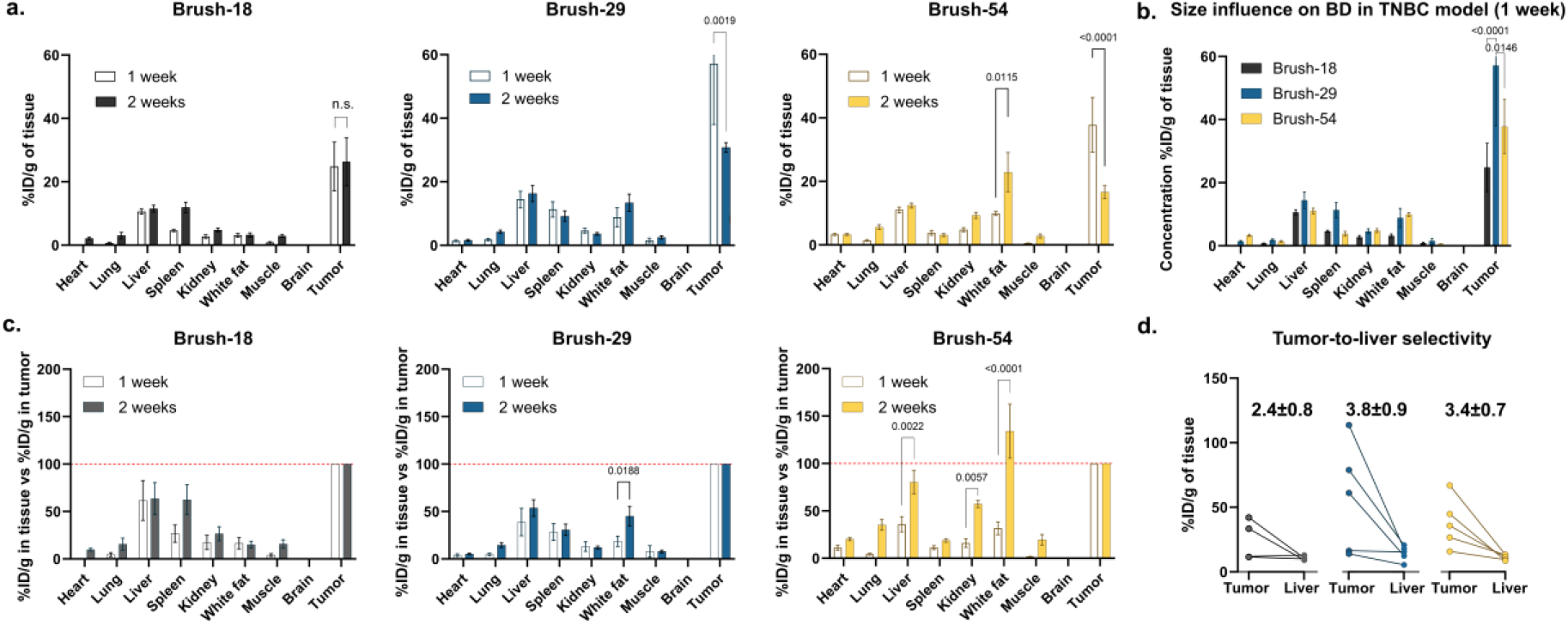
Biodistribution in 4T1 TNBC tumor-bearing mice 1 week after intraperitoneal administration. **a**, Comparison of biodistribution profiles for Brush-18, Brush-29 and Brush-54 at one (transparent bars)and two (filled bars) weeks after intraperitoneal administration (n=4-5 mice, average± SEM). **b**, Influence of Brush size on biodistribution in TNBC mice 1 week after intraperitoneal administration (n=4-5 mice, average± SEM). **c**, Influence of Brush size on accumulation in endogenous tissues versus accumulation in tumor at one (transparent bars) and two (filled bars) weeks after administration. Values are averages of tissue concentrations relative to concentrations in tumors, calculated individually for each mouse (n=4-5 mice, average± SEM). **d**, Comparison of Brush concentrations in tumor vs liver in individual mice at 7 days post administration. Values above the lines represent mean± SEM of the ratios.

To better assess biodistribution and tumor accumulation, we repeated the experiment, setting the end point at 7 days. Brush concentrations in all native tissues were mostly invariant between 7 and 14 day timepoints (**Fig. 4a**). At the 7-day mark, tumor concentrations of Brush-29 and Brush-54 were 1.9 and 2.3 times higher, respectively, compared to their concentrations at 14 days, confirming that ‘dilution’ of Brush concentration in tumors happened between days 7 and 14. Similar to 14-day timepoint results, the nanomaterial size had influence on tumor accumulation and Brush-29 was the most efficient one, reaching the mean value of 57.1 %ID/g of tumor, while Brush-18 and Brush-54 concentrations were respectively measured at 24.8 and 37.8 %ID/g. Remarkably, despite the tumor doubling in size between days 7 and 14 post-administration, the average concentrations of Brush-18 in the tumor were nearly equal at these timepoints. This implies that Brush-18 may still be biodistributing, possibly by migrating from other tissues.

In the absence of non-specific interactions and protein corona formation, the tumor accumulation is governed purely by the size of nanomaterial and it was a significant factor in tumor accumulation efficiency (**Fig. 4b**). Among three particles tested, Brush-29 was the most successful in passive accumulation in tumor. While both Brush-29 and Brush-18 are smaller than Brush-54 and must therefore more readily extravasate from capillaries and enter tumor, Brush-18 appears to also readily penetrate through tumor mass and return back to circulation. Thus, in the absence of receptor targeting or a mechanism that would facilitate retention of nanoparticles, the optimal D_H_ lies in the 18-54 nm range.

Recently, variations in vascular permeability to a 12 nm nanomaterial (genetically recombinant human ferritin nanocage) were shown both within a single tumor mass and among tumors of the same type, such as 4T1. Here, we observed that this variation influences the biodistribution of larger nanomaterials as well. Indeed, coefficient of variation (CoV) for Brush-18 concentration in tumors was 62% at day 7 and 65% at day 14 post administration. This clearly was due to properties of tumors and not technical limitations of the experiments or analysis, as CoV for liver was 4.1 times lower at only 15%. In cases of Brush-29 and Brush-54, CoV for concentrations in tumor were also high and equal to 75% and 51% at day 7. These results indicate, that although tumor is the major accumulation site due to the EPR effect, the efficiency of this accumulation varies greatly due to inherent heterogeneity in tumor physiology.

A primary benefit of using nanoparticles for drug delivery in cancer therapy is their potential to enhance the therapeutic index of chemotherapeutic agents through the selective accumulation in cancerous tissues. To assess the selectivity of accumulation in tumors, we normalized the concentrations observed in tissues to the concentration within the tumor for each individual animal in the TNBC groups (**Fig. 4c**). Based on the averages from the pairwise comparisons, we observed that the highest accumulation of Brushes was predominantly in tumors across all tested groups, with the exception of Brush-54 at 14 days post administration. The minimal accumulation of all Brushes in the heart compared to the tumor at 7-day time point is noteworthy. Considering the cardiotoxic risks associated with many chemotherapeutics, the pronounced tumor-to-heart selectivity that can be achieved through passive accumulation exemplifies the benefits of nanoparticle-based drug delivery in cancer therapy that can be achieved through the EPR effect.

Relative accumulation in other organs, particularly liver, was considerably more pronounced. Liver is another common site of nanoparticle accumulation and the tumor-to-liver concentration ratio is frequently used to gauge the specificity of nanoparticle delivery. The non-opsonizing properties and very long circulation times of our developed nanomaterials allowed us to evaluate the limits of this selectivity that can be achieved through EPR (**Fig. 4d**). Brush-29 exhibited the highest average accumulation in tumor and also demonstrated the best tumor-to-liver selectivity with an average value of 3.8. In comparison, Brush-18 and Brush-54 had selectivity of 2.4 and 3.4, respectively. These values are notably higher than what is conventionally observed with nanoparticles not designed to resist opsonization. However, there was a substantial variability in selectivity, which we attribute to the inherent heterogeneity of tumors. Indeed, for the best performing Brush-29, measured tumor-to-liver concentration ratio varied from 1.1 to 6.4.

## Conclusions

Leveraging the EPR effect has been a cornerstone in the development of nanoparticles for cancer diagnostics and therapy for decades. However, the true potential and limits of the EPR effect have remained elusive. In this study, we took the PEGylation approach to its extreme and created unimolecular nanoparticle surrogates, bottle brush polymers, which are almost entirely composed of PEG. The Brushes we developed do not opsonize, have exceptionally long circulation times(days), and are even able to navigate through lymphatic system as was evident from PK profiles upon *i*.*p*. administration. These attributes enabled us to investigate the achievable limits of tumor accumulation due to the EPR effect, eliminating confounding variables. While this study focused on commonly used 4T1 TNBC model, developed nanomaterials can undoubtedly serve as a tool to study limits of the EPR in other tumors.

Presence of 4T1 TNBC tumor had no impact on pharmacokinetic profiles and tumor did not act like a sink for nanomaterials. Counterintuitively, the circulation time decreased with an increase in Brush size, while uniformity of biodistribution increased. These findings emphasize the critical balance between endothelial permeability and the capacity for diffusion through, and subsequent exit from, a tissue. This balance was also evident in tumor accumulation. Cancerous tissue was the primary site of accumulation for all Brushes. Brush-29 demonstrated the highest accumulation, revealing that the optimal size (D_H_) for passive tumor accumulation via EPR lies between 18 and 54 nm.

Brush-29, while unlikely to correspond precisely to the optimal size for tumor accumulation, must be in close proximity to it. Thus, the upper average limit of tumor accumulation attributable to the EPR effect is proximate to 60 %ID/g, and the maximal tumor-to-liver selectivity approaching a value of 4. While these numbers are encouraging, tumor heterogeneity leads to broad variability in both. For Brush-29 tumor accumulation ranged from 14 to 114 %ID/g and tumor-to-liver selectivity from 1.1 to 6.4. This variability poses substantial challenges. It becomes evident, that it is impossible to develop nanoparticles that rely solely on the EPR effect to achieve uniform efficacy in tumor targeting not just across various types of cancer, but even within the same tumor type in different patients. Without active targeting or retention-promoting features, EPR-driven nanoparticles can only be successful through personalized medicine approach and would require testing for tumor predisposition for nanoparticle accumulation.

## METHODS

### Synthesis of labelled Brushes

64 uL of DMSO solution of AEMA (2.3 mg, 13 µmol) was placed into a 10 mL rbf flask containing 2kPEG-NMA (3.0g, 140 µmol) in a mixture of DMSO (1 mL) and 0.06M acetate buffer in water (7 mL). DMSO solution of 4-((((2-carboxyethyl)thio)carbonothioyl)thio)-4-cyanopentanoic acid (chain transfer agent, CTA, varied from 0.0005-0.0080 eq) and DMSO solution of 4,4’-Azobis(4-cyanovaleric acid) (initiator, V-501, 0.5 eq with respect to CTA). Reaction solution was deoxygenated with argons gas and polymerizations were carried out at 70 °C for 22h. Reaction mixture was cooled, exposed to air, diluted with 90 mL of DI water, dialyzed (MWCO 50kDa) against DI water and finally lyophilized. Obtained powder was dissolved in dichloromethane, precipitated with diethyl ether and dried. Product was dissolved in 5 mL of 0.06 M acetate buffer, V-501 (100 eq) were added, solution was deoxygenated and heated to 80 C for 10 hours. Reaction mixture was cooled, diluted with 90 mL of DI water, dialyzed (MWCO 50kDa) against DI water and lyophilized to yield bottle brush polymers. Fluorescent labelling: bottle brush polymers were dissolved in 0.1 M NaHCO_3_, and placed in an ice-bath. DMSO solution of AlexaFluor(AF) NHS ester (Brush-18: AF750; Brush-29: AF647; Brush-54: AF680) was added and reaction was allowed to proceed overnight. DMSO solution of MeO-PEG_12_-COONHS (100 eq) was to cap any remaining primary amines and reaction was stirred for 6h. Reaction mixture was dialyzed and washed with DI water using centrifugal concentrator (MWCO 100kDa) until there was no appreciable fluorescence in the filtrate. Labelled polymer solutions were filtered through 0.2 µm syringe filter and lyophilized to yield Brushes used further in the studies.

### DLS measurements

The lyophilized Brush powder was dissolved in PBS (5 mg/mL) and filtered through a sterile 0.2 μm PES membrane. DLS measurements were conducted at 37 °C in disposable microcuvettes using Zetasizer Nano (Malvern Panalytical). Data was analysed using Zetasizer software (version 8.02), extracted and plotted with GraphPad Prism software (version 10.0.1).

### FCS measurements

Brush-18, Brush-29 and Brush-54 were dissolved in 0.2 μm-filtered Balb/c mouse serum prepared in-house at 10, 29 and 90 μg/mL concentrations, correspondingly. Sample aliquots were immediately placed on custom-made flat glass slides with parallel channels (approx. 8 μL volume), sealed and subjected to FCS measurements. The remaining solutions in serum were sealed and incubated in the dark at 37 °C while shaking at 250 rpm. Aliquots from incubated samples were taken after 24 h, 3 days, 5 days and 7 days and for FCS measurements. FCS was performed using a Leica Stellaris 8 inverted confocal microscope (Leica Microsystems GmbH) with a HC PL APO CS2 86x/1.20 water immersion objective. Excitation laser lines of 653 nm, 681 nm and 688 nm were delivered by the Stellaris supercontinuum pulsed (80 MHz) white light laser, each line was tuned with Stellaris acousto-optical beam splitter. GaAsP power hybrid detectors HyD X or HyD R were used for photon counting in the red and far-red emission spectra, correspondingly. Autocorrelation data were acquired using Leica built-in autocorrelator, data were extracted and plotted using GraphPad Prism software (version 10.0.1).

### Healthy mice and TNBC models

All animal experiments Experimental procedures were performed in accordance with the guidelines reported in the EU Directive 2010/63/EU and local laws and policies. All performed procedure were approved by the Latvian Animal Protection Ethical Committee of the Food and Veterinary Service (Riga, Latvia). Balb/c mice (4-6 weeks old) were obtained from the Laboratory Animal Centre, University of Tartu (Tartu, Estonia) and were used in experiments when reached 8 weeks of age. Mice were housed in individually ventilated cages, 5 animals per cage, with unlimited access to food and water. Mice were maintained under standard housing conditions with temperature range of 21–23 °C, 12-hour light and dark cycle and relative humidity level of 50 ± 5%. Before the experiment, the mice were adapted to housing conditions for more than one week. For the orthotopic breast cancer model, Balb/c female mice were orthotopically inoculated with 4T1 cell suspension (10^5^ 4T1 cells per mouse) in the fourth inguinal mammary fat pad (60 μL/mouse). Tumor volume was measured with a caliper every other day and calculated using the formula: (width^2^×length)/2. Seven days after inoculation tumor-bearing mice were assigned to groups to ensure uniformity in average tumor volume and distribution across the groups, after which nanomaterial administration commenced. Each group in both healthy and TNBC cohorts consisted of 6 animals. Throughout the duration of study animals were monitored for clinical signs necessitating immediate intervention and immediate humane termination^29^.

### Pharmacokinetic and Biodistribution studies

Brush solutions in PBS were prepared at equimolar concentrations for Brush-18, Brush-29 and Brush-54 (100, 290 and 900 μg/mL, respectively) and sterilized by filtration through 0.2 μm PES syringe filters. Each mouse received 100 μL bolus injection of Brush solution either via tail vein (healthy i.v. groups) or into the lower right quadrant of the abdomen (healthy i.p. and TNBC groups). We first established that Brushes do not associate with blood cells and persist in the serum following separation, and used whole blood for the quantification of Brush concentrations minimizing additional manipulation prior to quantitation. Blood samples (10-20 μL) were collected one day prior to administration and 1h, 4h, 8h, 1d, 3d, 7d, 14d (i.v. groups) or 0.5h, 2h, 6h, 18h, 43h, 4d, 7d, 14d (i.p. and TNBC groups) after administration, weighted and frozen. Mice were anaesthetized with 4% isoflurane, transcardially perfused with PBS (3 ml/min for 5 min) seven days (1-week TNBC groups) or 14 days (healthy groups and 2-week TNBC groups) after Brush administration. Tissues were collected and frozen. Blood samples were thawed, diluted with 1% Triton-X 100 solution in PBS at 1:9 (w/v) ratio, and incubated for 20 minutes at room temperature while agitating at 350 rpm. 100 µL of each sample were transferred to 96-well plate and fluorescence spectra were obtained using the Tecan Infinite M1000 microplate reader. Organs and tissues were homogenized in PBS at 1:4 (w/v) ratio using stainless steel beads and a beadmill (OMNI International Inc). 100 µL of each sample were transferred to 96-well plate and fluorescence spectra were obtained using the Tecan Infinite M1000 microplate reader. Standard curves were generated by measuring fluorescence of homogenized blood and tissues samples from untreated animals, which were spiked with known concentrations of Brushes and used to calculated Brush concentrations (w/w) and expressed as %ID/g of tissue after normalization to the injected dose

### PK modelling

Brush concentrations in blood were fitted into two-compartmental model using PKSolver add-in^30^. Measured blood concentrations and obtained PK curves were plotted using GraphPad Prism software (version 10.0.1).

### Statistical analysis

Statistical analysis was done using GraphPad Prism software (version 10.0.1). The statistical differences were analyzed using one-way/two-way ANOVA, for two and three groups comparison respectively, followed by Šídák’s (matching tissue Brush concentration between two groups), Tukey’s (matching tissues Brush concentration between three groups and tissue-to-tissue Brush concentration comparison within a single group), and Dunnett’s (Brush concentration comparison between endogenous tissues and tumor) multiple comparisons tests. All the results are expressed as mean ± s.e.m., unless noted otherwise.

## Data availability

The authors declare that data supporting the findings of this study are available within the article and its Supplementary Information. All relevant data can be made available upon reasonable request to the corresponding author.

## Author contributions

R.S. collected data, conducted data analysis and performed all the non *in vivo* experiments. B.S. collected data and performed all *in vivo* experiments. I.C. performed FCS experiments and data analysis. G.S. performed in vivo experiments. A.S. conceived the idea, conducted data analysis, designed and supervised all studies and wrote the manuscript. All authors edited the manuscript.

## Acknowledgements

This work was supported by the Latvian Council of Science grants No. lzp-2020/2-0048, lzp-2021/1-0593 and EU Horizon 2020 research and innovation programme grant agreement No. 857287.

## Notes

### Competing Interest Statement

The authors have declared no competing interest.

## References

1. Maeda, H., Matsumoto, T., Konno, T., Iwai, K. & Ueda, M. Tailor-Making of Protein Drugs by Polymer Conjugation for Tumor Targeting: A Brief Review on Smancs. Journal of Protein Chernistry vol. 3 (1984).

2. Matsumura, Y. & Maeda2, H. A New Concept for Macromolecular Therapeutics in Cancer Chemotherapy: Mechanism of Tumoritropic Accumulation of Proteins and the Antitumor Agent Smancs1. http://aacrjournals.org/cancerres/article-pdf/46/12_Part_1/6387/2424185/cr04612p16387.pdf (1986).

3. Fang, J., Islam, W. & Maeda, H. Exploiting the dynamics of the EPR effect and strategies to improve the therapeutic effects of nanomedicines by using EPR effect enhancers. Advanced Drug Delivery Reviews vol. 157 142–160 Preprint at 10.1016/j.addr.2020.06.005 (2020).

4. Sindhwani, S. et al. The entry of nanoparticles into solid tumours. Nat Mater 19, 566–575 (2020).

5. Zhu, M. et al. Machine-learning-assisted single-vessel analysis of nanoparticle permeability in tumour vasculatures. Nat Nanotechnol 18, 657–666 (2023).

6. de Lázaro, I. & Mooney, D. J. Obstacles and opportunities in a forward vision for cancer nanomedicine. Nature Materials vol. 20 1469–1479 Preprint at 10.1038/s41563-021-01047-7 (2021).

7. Ouyang, B. et al. The dose threshold for nanoparticle tumour delivery. Nat Mater 19, 1362–1371 (2020).

8. Chen, S. et al. Enhanced tumour penetration and prolonged circulation in blood of polyzwitterion–drug conjugates with cell-membrane affinity. Nat Biomed Eng 5, 1019–1037 (2021).

9. Mishra, P., Nayak, B. & Dey, R. K. PEGylation in anti-cancer therapy: An overview. Asian Journal of Pharmaceutical Sciences vol. 11 337–348 Preprint at 10.1016/j.ajps.2015.08.011 (2016).

10. Sosale, N. G., Spinler, K. R., Alvey, C. & Discher, D. E. Macrophage engulfment of a cell or nanoparticle is regulated by unavoidable opsonization, a species-specific ‘Marker of Self’ CD47, and target physical properties. Current Opinion in Immunology vol. 35 107–112 Preprint at 10.1016/j.coi.2015.06.013 (2015).

11. Vu, V. P. et al. Immunoglobulin deposition on biomolecule corona determines complement opsonization efficiency of preclinical and clinical nanoparticles. Nat Nanotechnol 14, 260–268 (2019).

12. Moein Moghimi, S., Simberg, D., Skotland, T., Yaghmur, A. & Christy Hunter, A. The interplay between blood proteins, complement, and macrophages on nanomedicine performance and responses. Journal of Pharmacology and Experimental Therapeutics vol. 370 581–592 Preprint at 10.1124/jpet.119.258012 (2019).

13. Kang, H. et al. Size-Dependent EPR Effect of Polymeric Nanoparticles on Tumor Targeting. Adv Healthc Mater 9, (2020).

14. Yadav, D. & Dewangan, H. K. PEGYLATION: an important approach for novel drug delivery system. Journal of Biomaterials Science, Polymer Edition vol. 32 266–280 Preprint at 10.1080/09205063.2020.1825304 (2021).

15. Suk, J. S., Xu, Q., Kim, N., Hanes, J. & Ensign, L. M. PEGylation as a strategy for improving nanoparticle-based drug and gene delivery. Advanced Drug Delivery Reviews vol. 99 28–51 Preprint at 10.1016/j.addr.2015.09.012 (2016).

16. Schmidt, M. M. & Wittrup, K. D. A modeling analysis of the effects of molecular size and binding affinity on tumor targeting. Mol Cancer Ther 8, 2861–2871 (2009).

17. Sykes, E. A., Chen, J., Zheng, G. & Chan, W. C. W. Investigating the impact of nanoparticle size on active and passive tumor targeting efficiency. ACS Nano 8, 5696–5706 (2014).

18. Ries, J. & Schwille, P. Fluorescence correlation spectroscopy. BioEssays 34, 361–368 (2012).

19. Wang, H., Lin, Y., Nienhaus, K. & Nienhaus, G. U. The protein corona on nanoparticles as viewed from a nanoparticle-sizing perspective. Wiley Interdiscip Rev Nanomed Nanobiotechnol 10, (2018).

20. Leypoldt, J. K. Solute Transport Across the Peritoneal Membrane. Journal of the American Society of Nephrology 13, S84–S91 (2002).

21. Deplaine, G. et al. The sensing of poorly deformable red blood cells by the human spleen can be mimicked in vitro. Blood 117, e88–e95 (2011).

22. Zapotoczny, B., Szafranska, K., Kus, E., Chlopicki, S. & Szymonski, M. Quantification of fenestrations in liver sinusoidal endothelial cells by atomic force microscopy. Micron 101, 48–53 (2017).

23. Arroyo-Crespo, J. J. et al. Characterization of triple-negative breast cancer preclinical models provides functional evidence of metastatic progression. Int J Cancer 145, 2267–2281 (2019).

24. Malekian, S. et al. Expression of diverse angiogenesis factor in different stages of the 4T1 tumor as a mouse model of triple-negative breast cancer. Adv Pharm Bull 10, 323–328 (2020).

25. Schrörs, B. et al. Multi-Omics Characterization of the 4T1 Murine Mammary Gland Tumor Model. Front Oncol 10, (2020).

26. Madera, L., Greenshields, A., Coombs, M. R. P. & Hoskin, D. W. 4T1 murine mammary carcinoma cells enhance macrophage-mediated innate inflammatory responses. PLoS One 10, (2015).

27. Walker, W. H. et al. Mammary Tumors Induce Central Pro-inflammatory Cytokine Expression, but Not Behavioral Deficits in Balb/C Mice. Sci Rep 7, (2017).

28. Steenbrugge, J., de Jaeghere, E. A., Meyer, E., Denys, H. & de Wever, O. Splenic Hematopoietic and Stromal Cells in Cancer Progression. Cancer Res 81, 27–35 (2021).

29. Workman, P. et al. Guidelines for the welfare and use of animals in cancer research. British Journal of Cancer vol. 102 1555–1577 Preprint at 10.1038/sj.bjc.6605642 (2010).

30. Zhang, Y., Huo, M., Zhou, J. & Xie, S. PKSolver: An add-in program for pharmacokinetic and pharmacodynamic data analysis in Microsoft Excel. Comput Methods Programs Biomed 99, 306–314 (2010).

