## Supplemental figures 1-4 for "Exploring the Limits of EPR-driven Tumor Accumulation with Non-opsonizing Nanomaterials"

### Supplementary information

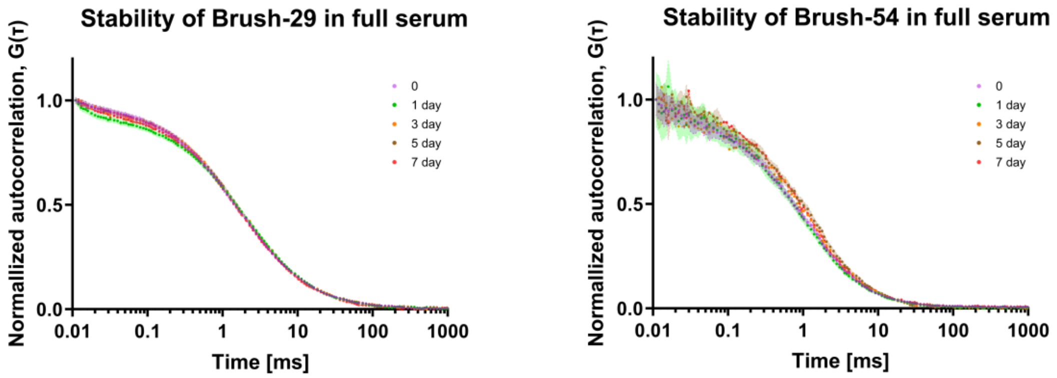


**Supplementary Fig. 1 | Resistance to opsonization in full serum.** Fluorescence correlation spectroscopy autocorrelation curves did not change during the 7 day period of incubation in full mouse serum at 37 °C, indicating there were no changes in Brush diffusion and no opsonization or protein corona formation.

| Results of Šídák's multiple comparisons test | | | |
|---|---|---|---|
| Time, d | Adjusted P Value, healthy (i.p.) vs 4T1 (i.p.) | | |
| | Brush-18 | Brush-29 | Brush-54 |
| 0.02 | 0.8241 | 0.9589 | 0.7441 |
| 0.08 | <0.0001 | 0.0012 | 0.9963 |
| 0.25 | 0.1588 | 0.7448 | 0.9997 |
| T0.75 | >0.9999 | 0.9995 | 0.8161 |
| 1.79 | 0.9982 | >0.9999 | 0.2053 |
| 4.00 | 0.971 | >0.9999 | 0.8444 |
| 7.00 | 0.9864 | >0.9999 | >0.9999 |
| 14.00 | >0.9999 | >0.9999 | >0.9999 |

**Supplementary Fig. 2 |** p-values for comparison of Brush concentrations measured in blood of healthy mice vs. Brush concentrations measured in blood of 4T1 tumor-bearing mice at matching time points.

| | p-values of Dunnett's multiple comparisons test | | | | | |
|---|---|---|---|---|---|---|
| | 1-week post administration | | | 2-weeks post administration | | |
| | Brush-18 | Brush-29 | Brush-54 | Brush-18 | Brush-29 | Brush-54 |
| Tumor vs. Heart | <0.0001 | <0.0001 | <0.0001 | <0.0001 | <0.0001 | 0.0012 |
| Tumor vs. Lung | <0.0001 | <0.0001 | <0.0001 | <0.0001 | <0.0001 | 0.009 |
| Tumor vs. Liver | 0.0043 | 0.0004 | <0.0001 | 0.0028 | <0.0001 | 0.6745 |
| Tumor vs. Spleen | <0.0001 | 0.0001 | <0.0001 | 0.0037 | <0.0001 | 0.001 |
| Tumor vs. Kidney | <0.0001 | <0.0001 | <0.0001 | <0.0001 | <0.0001 | 0.1428 |
| Tumor vs. White fat | <0.0001 | <0.0001 | <0.0001 | <0.0001 | <0.0001 | 0.2594 |
| Tumor vs. Muscle | <0.0001 | <0.0001 | <0.0001 | <0.0001 | <0.0001 | 0.0008 |
| Tumor vs. Brain | <0.0001 | <0.0001 | <0.0001 | <0.0001 | <0.0001 | <0.0001 |

**Supplementary Fig. 3 |** Tumor was the primary site of accumulation of all tested Brushes. p-values of Dunnett's multiple comparisons test performed within each TNBC group following a one-way ANOVA between Brush concentration in tumor vs Brush concentration in other tissues.

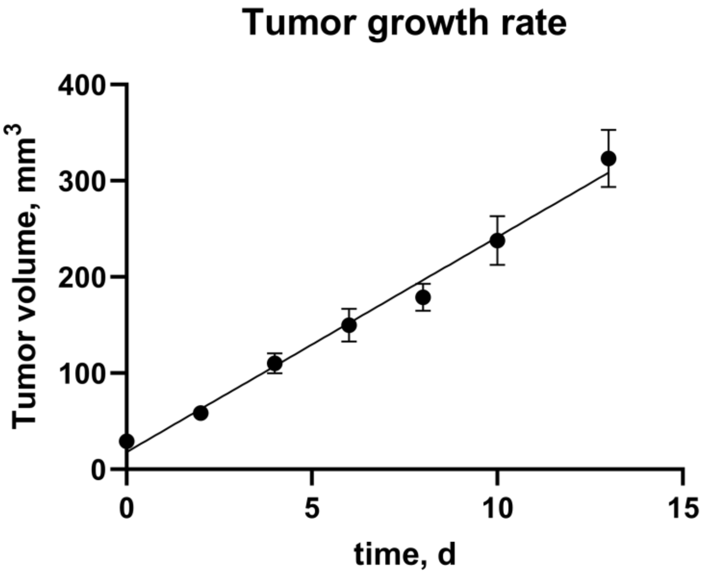


**Supplementary Fig. 4 |** Growth rate of 4T1 tumors observed in 2-week TNBC groups. Dimensions were measured with a caliper and volumes calculated using the formula: (width$^2$×length)/2. Data points represent mean±SEM (n=15).
